# Gene-centered representation of coding and regulatory variation enables outcome prediction

**DOI:** 10.64898/2026.01.27.701808

**Authors:** Ksenia Sokolova, Vessela N Kristensen, Christopher Y Park, Chandra Theesfeld, Laura Mariani, Matthias Kretzler, Olga G. Troyanskaya, Cure Glomerulonephropathy (CureGN) Study Consortium

**Author notes:** A list of CureGN Consortium members and affiliations appears in the Supplementary Information.

## Abstract

Integrating coding and regulatory variation into unified, interpretable representations remains a challenge in functional genomics. Current approaches either focus on common variants or analyze individual variants in isolation, missing the cumulative, cell-type-specific impact of both coding and noncoding variants on each gene. We present Volaria, a computational framework that integrates coding and regulatory genetic variation into unified, gene-centered representations for disease outcome prediction from whole-genome sequencing. Volaria leverages deep learning models to capture variant effects on cell-type-specific gene expression and integrates them with AI-predicted exonic variant pathogenicity to produce representations that capture the cumulative effect of genome-wide rare and common variation. Applied to whole genomes of individuals with rare glomerular diseases, Volaria predicts individual outcomes directly from germline sequence, demonstrating that structured, cell-type-aware representations capture predictive signals beyond population-based polygenic risk scores and unstructured representations. Importantly, the framework identifies context-specific biological mechanisms, providing interpretability that can be aligned with clinical measurements. By encoding genome-wide variation into compact and biologically grounded representations, Volaria provides a scalable foundation for genome interpretation and individualized outcome modeling from germline sequence, complementing phenotypic and clinical information in the future integrative frameworks.

## Introduction

Leveraging whole-genome sequencing (WGS) data to connect genetic variation to clinical outcomes remains a central challenge in genomic medicine. WGS has led to important insights into common and rare variants associated with phenotypes [1], delivered diagnoses for rare disease [2], and led to discoveries in pharmacogenomics and drug selection [3,4]. However, although WGS captures rare and common variants with base-pair resolution across coding and non-coding regions, existing analysis approaches fail to effectively integrate this information and convert it into actionable insights about disease progression and treatment.

A popular strategy to quantify genetic susceptibility to complex diseases is to use Polygenic Scores (PGS) [5] that act by aggregating the effects of multiple common variants into a single predictive score. While widely adopted in research and in clinical risk modeling, PGS models rely on GWAS-derived associations and inherit their limitations, including focus on common variants and lack of tissue- and cell-type-specific resolution. Most importantly, PGS primarily capture disease susceptibility in healthy individuals, rather than outcomes such as disease progression, treatment response, or cell-type specific mechanisms within affected individuals.

Many other approaches focus on coding variation, often prioritizing common protein-altering variants based on constraint or allele frequency [6,7]. While informative in some settings, these models exclude regulatory variation by design and assume relatively direct, high-penetrance effects. As a result, they miss contributions of non-coding variation and the cumulative impact of many individually weak signals.

These limitations are especially pronounced in the context of complex diseases, where rare and common variants distributed across coding and regulatory regions collectively influence pathogenesis [8–11]. The high dimensionality of WGS data compounds this challenge, as each individual carries a variable number of variants spread throughout the genome, with widely differing functional impact depending on tissue, cell type and regulatory context. Recent machine learning approaches have advanced prediction of biological variant impact at multiple regulatory levels. Deep learning models [12–15], including convolutional and transformer-based architectures, have significantly improved the interpretability of genomic variation, particularly in non-coding regions. However, these methods operate at the variant level, lacking the ability to model the cumulative impacts of variants across the genome. There is a need for a structured, interpretable, individual-level representation of genome-wide variation that integrates these diverse signals.

We introduce Volaria, a computational framework that integrates both exonic and regulatory variants into a single, biologically informed, explainable embedding for each individual. Volaria incorporates all variants, common and rare, coding and noncoding, without prior assumptions such as allele frequency thresholds or variant-level significance. Volaria leverages cell-type-specific regulatory effect predictions [14] and functional assessments of exonic variation [15] to generate unified gene-centered features, which are aggregated into individual-level embeddings. We refer to these gene-centered feature matrices as ‘embeddings’ to emphasize their fixed dimensionality and reusability across prediction tasks. By unifying these effects, Volaria generates biologically meaningful context-specific embeddings that are both scalable and interpretable across diverse disease contexts.

We demonstrate Volaria’s utility for predicting clinical outcomes from only genome sequences in individuals with rare kidney glomerular diseases from the Cure Glomerulonephropathy study (CureGN) [16]. These conditions are characterized by substantial clinical and genetic heterogeneity, and a subset of affected individuals will progress to kidney failure. Critically, Volaria enables outcome modeling from germline WGS alone, information available at or before diagnosis, without requiring clinical covariates or longitudinal measurements. We further leverage the interpretability of the framework to identify specific genes underlying outcome predictions and connect them to clinical readouts. Together, Volaria’s gene-centered, cell type-aware representations provide a scalable foundation for genome-based outcome modeling and mechanistic insight, complementary to clinical data and suitable for future integrative precision health approaches.

## Results

Volaria constructs cell-type-specific, gene-centered representations of an individual’s genome by integrating the predicted effects of both coding and non-coding variants (Figure 1A). These are aggregated into fixed-length individual-level embeddings used as inputs to downstream prediction models (Figure 1B). While the full system is referred to as Volaria, its modular design allows embeddings to be flexibly applied across diverse predictive tasks. Both regulatory effects, which capture cis-regulatory influences on cell-type-specific gene expression, and exonic pathogenicity effects are predicted directly from individual genomic sequence [14][15]. All variants are retained and aggregated at the gene level for each individual. These embeddings map the high-dimensional space of an individual’s genomic variation into unified, gene-centered representations that reflect cell-type and tissue-specific molecular perturbations (Figure 1A; Methods).

**Figure 1.**
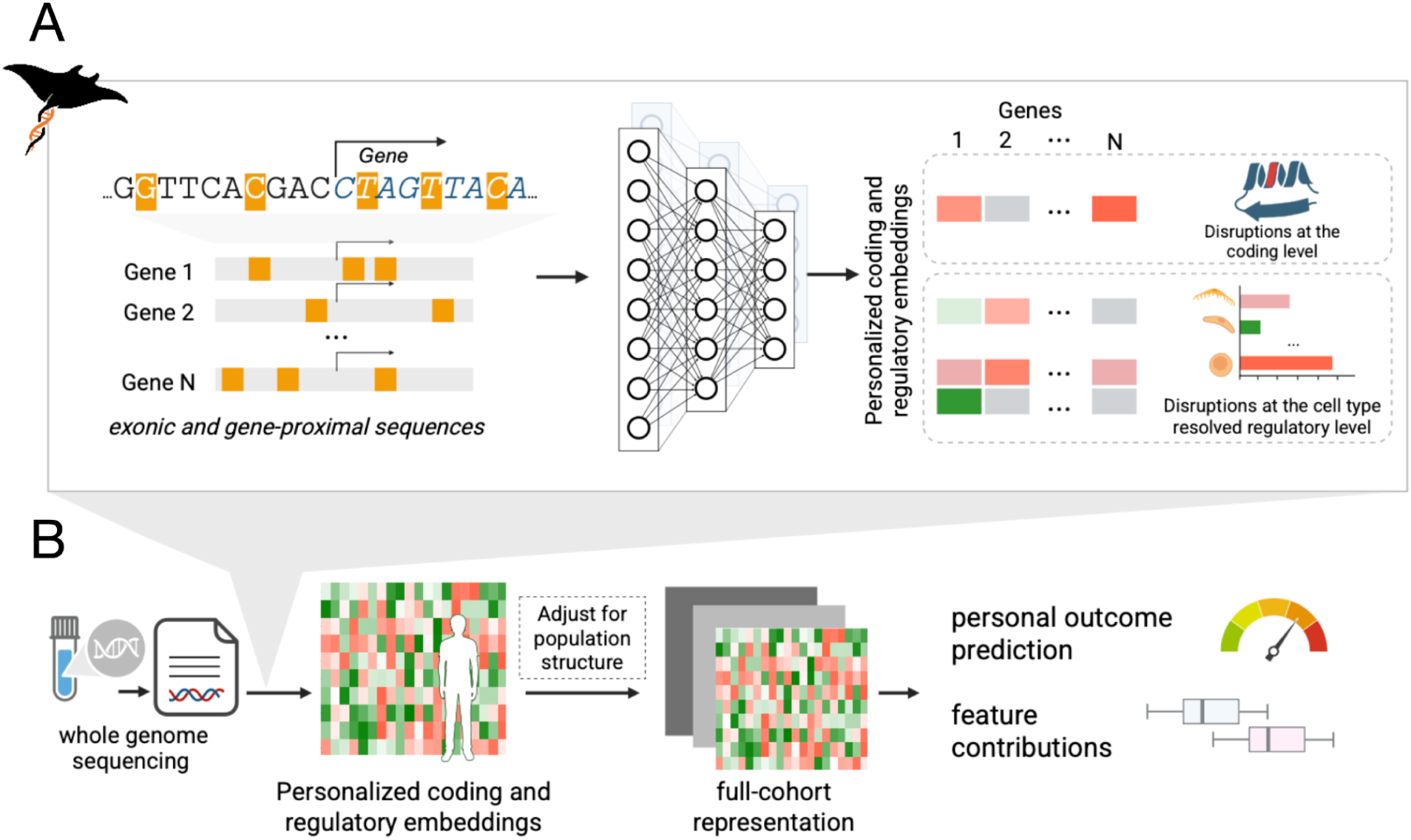
Overview of the Volaria framework. (A) Volaria integrates the effects of both coding and regulatory variants across the genome to produce gene-centered functional representations. Coding variants are evaluated using protein impact models, while non-coding variants are scored for predicted disruption of gene expression across six kidney cell types, including kidney immune cell types. To enable comprehensive modeling of the functional impact of genetic variation, no allele frequency filtering is applied (standard genotype QC used, Methods). (B) For each individual, Volaria processes WGS data to generate a matrix of gene-level variant effects (embeddings). These matrices have consistent dimensionality across individuals and serve as inputs for downstream predictive and exploratory analyses. Prior to cohort integration, to account for population structure, this matrix is adjusted by regressing out the top principal components (Methods). The resulting patient embeddings support multiple downstream applications, including interpretable outcome prediction.

### Volaria predicts clinical outcomes from genome sequence alone

We applied Volaria to WGS data from the Cure Glomerulonephropathy (CureGN [16]) study, a longitudinal clinical study of individuals with four rare glomerular diseases, to predict disease progression and treatment response (Figure 1B). CureGN provides matched WGS, deep clinical phenotyping, and nearly a decade of follow-up clinical outcomes data. We used the GTEx WGS dataset [17], which included individuals with kidney disease, as an external validation cohort. We further benchmarked Volaria against polygenic scores (PGS) for kidney failure from previously published studies.

Through consultation with nephrologists, we selected three clinically meaningful outcomes to predict: (1) 40% decline in estimated glomerular filtration rate (eGFR), a clinically meaningful threshold for predicting future kidney failure and mortality; (2) kidney failure, a condition where kidneys have permanently lost their ability to filter waste and excess fluid from the body with severe quality of life consequences, and (3) steroid resistance, the state under which kidneys do not respond to treatment with corticosteroids. Kidney failure and 40% eGFR decline are time-resolved endpoints, while steroid resistance was treated as a binary label independent of time (Figure 2A). Among 1,846 individuals in the cohort, 267 reached kidney failure, 329 experienced a 40% eGFR decline (median times of 687 and 775 days post-enrollment), and steroid resistance affected 199 individuals (Figure 2B). To evaluate the informativeness of the Volaria embeddings, we trained regression models with these outcome-specific targets. Model parameters were optimized via cross-validation, and individuals without the event were included only if they had at least 2,000 days of follow-up (Methods). These outcomes allow us to evaluate Volaria’s ability to capture both progressive decline and treatment response.

**Figure 2.**
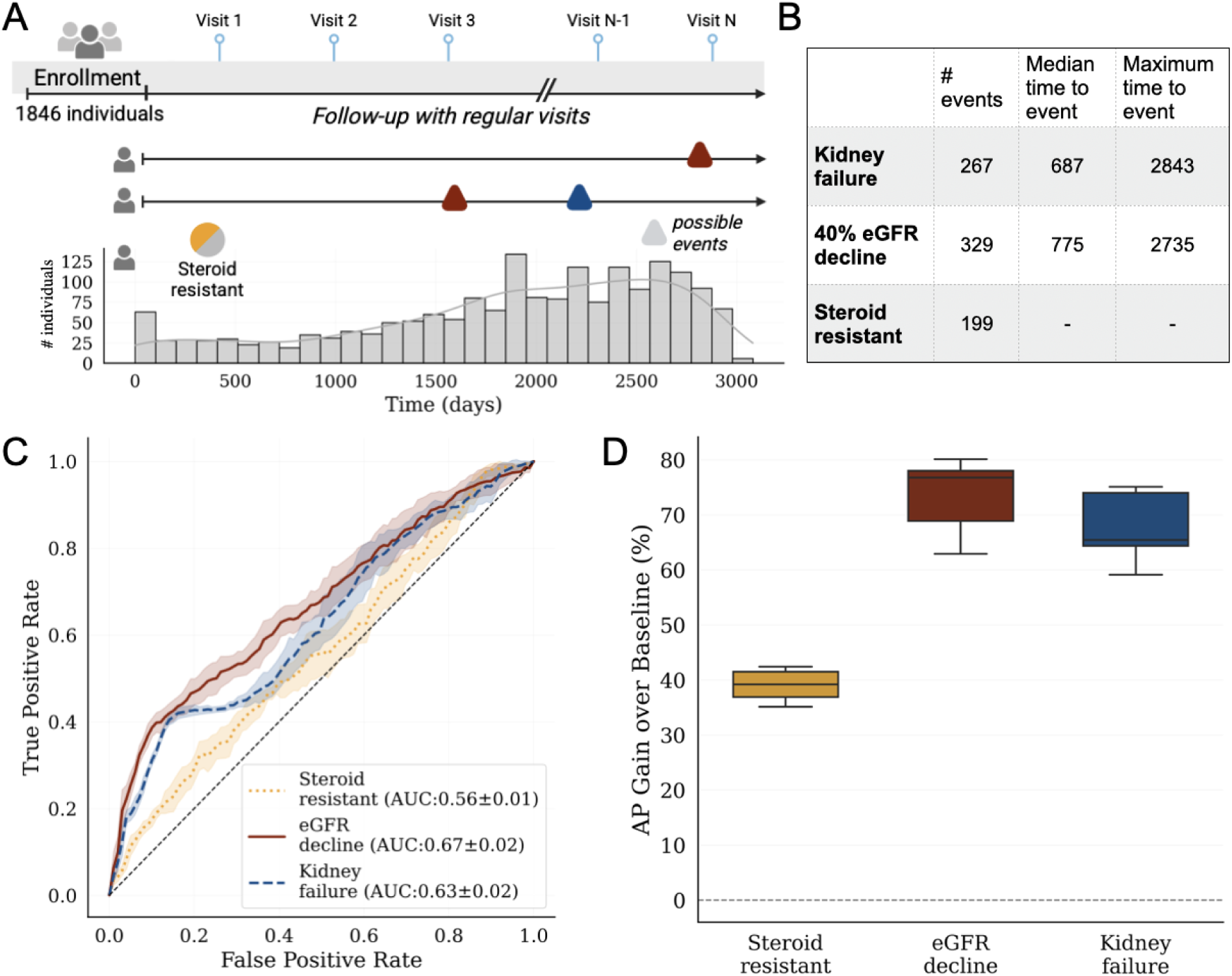
Evaluation of Volaria’s predictive performance across renal outcomes. (A) CureGN cohort overview: 1,846 individuals with WGS and longitudinal follow-up (up to 3,000 days). Three clinical outcomes were evaluated: 40% decline in estimated glomerular filtration rate (eGFR), progression to kidney failure, and steroid resistance. The first two outcomes are time-resolved events with known onset dates, while the latter is a status-based property without precise timing. Outcomes were not mutually exclusive. (B) Outcome frequencies and event timing. The 40% eGFR decline was the most common (329 individuals), followed by kidney failure, and steroid resistance. (C) Receiver operating characteristic (ROC) curves for genome-based outcome prediction on the test set. Prediction of 40% eGFR decline achieved the highest AUC (0.67), followed by kidney failure, and steroid resistance. Separate models were trained for each outcome using the same test set; error intervals denote standard deviation across random seeds. (D) Percent improvement in average precision (AP) over baseline for each outcome. All models substantially outperformed the baselines.

Volaria achieved meaningful predictive performance using only genomic information, without any clinical covariates, treatment history, or longitudinal data. Because the input is only germline WGS, predictions are available prior to diagnosis or before clinical data accumulates, making them complementary to dynamic clinical data. The observed discrimination for both kidney failure and 40% eGFR decline is notable given that genome sequence represents a fixed, pre-disease measurement (ROC AUC = 0.63 ± 0.02 for kidney failure; 0.67 ± 0.02 for 40% eGFR decline; Figure 2C, 2D, Supp. Table 1). Genome-based steroid resistance prediction was also possible but more challenging (ROC AUC 0.56 ± 0.01), likely reflecting the complex dependence of treatment response on non-genetic factors including treatment timing, protocol, and concurrent immunosuppression that are not captured by genetic sequence alone. These results demonstrate that Volaria’s gene-centered, cell type-aware genomic representations encode outcome-relevant signals even in the absence of clinical features, providing a complementary perspective to phenotypic risk modeling.

### Embedding structure contributes to the predictive signal

To evaluate whether gene- and cell-type-specific embeddings provide predictive value beyond unstructured representations, we compared Volaria to a series of embeddings with simplified structures. Specifically, we constructed flat functional embeddings for the same genes and variant scores by removing gene and cell-type-specific structure and aggregating variant scores by taking the maximum across the dropped dimensions (Methods). To isolate contributions from each signal type, we evaluated models using only regulatory predictions (Volaria flat regulatory), only exonic predictions (Volaria flat exonic), or providing both regulatory and exonic flat embeddings. Across all three renal outcomes, full Volaria embedding consistently outperformed unstructured ones (Figure 3A-C, Supp. Figure 1), indicating that gene-centered, cell-type-aware representations capture predictive signals not recoverable from unstructured inputs. Notably, the performance of simplified embeddings varied by outcome: regulatory signals performed comparably or better than exonic scores in predicting kidney failure and eGFR decline (Figure 3A-B), but were less informative for predicting steroid resistance (Figure 3C), suggesting task-specific relevance of variant classes.

**Figure 3.**
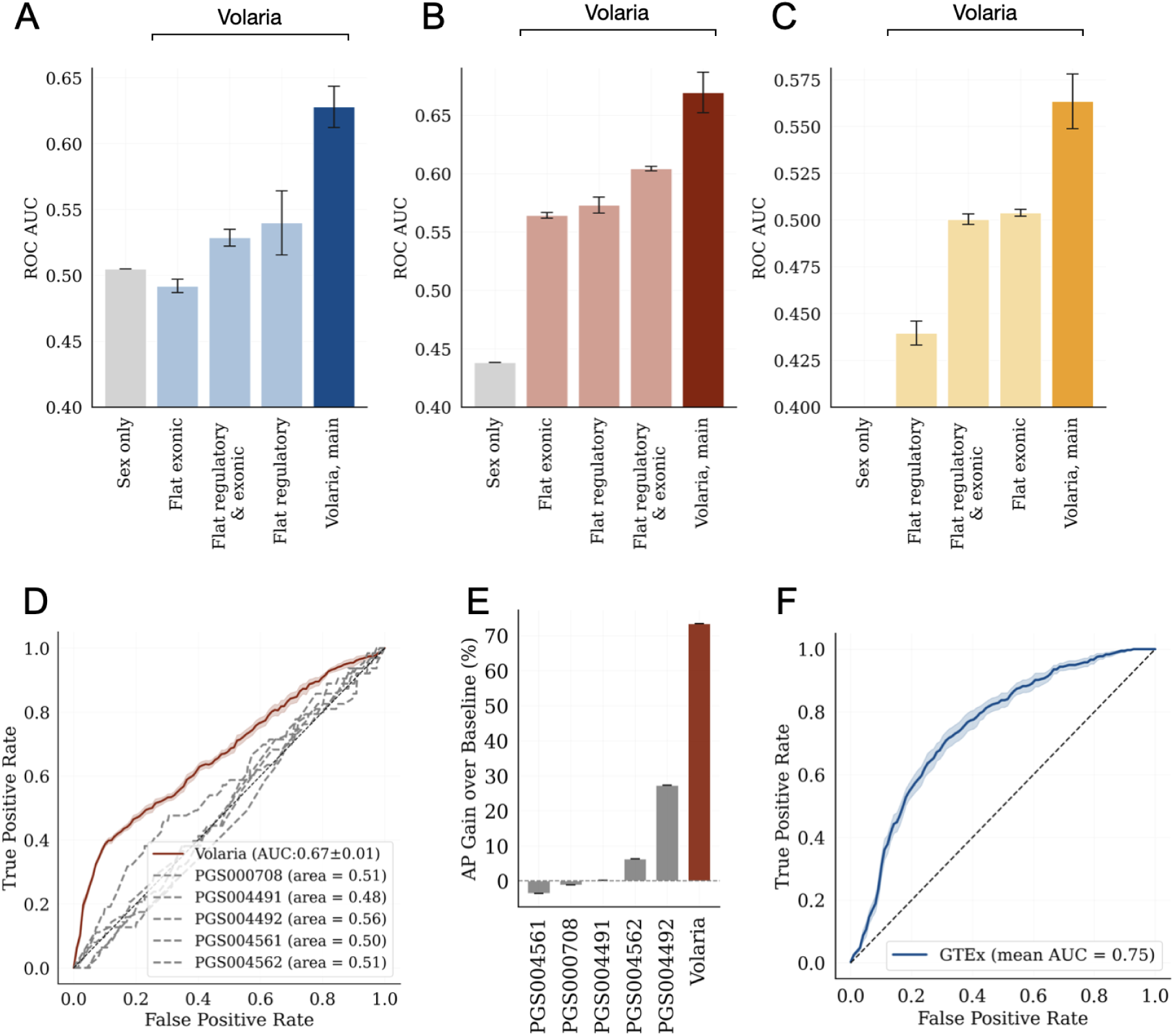
Extended performance evaluation. (A-C) Volaria’s embedding structure contributes to the predictive performance: (A) kidney failure, (B) eGFR decline, and (C) steroid resistance. To evaluate contribution of the gene-level resolution and cell-type, we compare flattened functional embeddings lacking gene-level or cell-type structure, separated into regulatory-only, exonic-only, and combined input configurations, as well as a simple baseline of only sex as an input. Volaria’s full structured embedding outperforms all non-structured variations, showing that both gene resolution and cell type specificity contribute to the downstream performance. (D) ROC AUC curves for predicting 40% eGFR decline using Volaria versus using polygenic scores (PGS) for kidney failure from the PGS Catalog; (E) Average precision improvements over baseline for the 40% eGFR decline using Volaria or each of the PGS scores. (F) GTEx cohort evaluation of Volaria models trained on CureGN (ROC AUC for kidney failure prediction in GTEx individuals). Volaria generalizes effectively across cohorts with different contexts and data collection protocols.

### Comparison to genome-wide polygenic risk models

To benchmark Volaria against established approaches for genome-wide risk prediction, we compared its performance to published polygenic scores for kidney-related traits. Specifically, we evaluated all five polygenic scores (PGS) associated with kidney failure or related phenotypes (acute and chronic renal failure) in the PGS Catalog: PGS000708 [18]; PGS004491, PGS004492, PGS004561, and PGS004562 [19]. These scores differ in phenotype definitions and modeling strategies. For example, PGS000708 defines kidney disease based on a combination of self-reported diagnoses and ICD-10 codes (N17–N19) in the UK Biobank, including both acute and chronic renal failure. At the same time, [19] did not perform aggregation over acute and chronic renal failure and provide different ways to compute scores (PGS004491 and PGS004492 represent acute and chronic renal failure, respectively, using LDpred2-based scores, while PGS004561 and PGS004562 represent acute and chronic renal failure, respectively and use elastic net regression, integrating disease-specific and risk factor polygenic scores). In accordance with these previous studies we computed PGS values for each of the five scores for each individual in the CureGN cohort. We then trained the same outcome-specific classifiers as used for Volaria, with identical train/test splits, for both 40% eGFR decline and kidney failure outcomes (Figure 3D–E, Supp. Figure 2).

Across all five scores and both outcomes, Volaria consistently outperformed PGS-based models in both ROC AUC and precision-recall metrics. For 40% eGFR decline, PGS average performance was 0.51±0.03 and the best-performing PGS (PGS004492, chronic renal failure LDpred2 [19]) achieved an ROC AUC of 0.56, substantially lower than Volaria’s 0.67. Volaria’s corresponding AP was more than twice that of the best-performing PGS model (Figure 3E). A similar pattern was observed for kidney failure, where PGS average performance was 0.56±0.03, with best performing PGS-based ROC AUC of 0.59, short of Volaria’s ROC AUC of 0.63, and with Volaria also achieving a two-fold improvement in PR AUC vs the best PGS (Supp.Figure 2).

### Volaria captures broad signal contributions

A subset of the CureGN participants have been previously reported to present with monogenic forms of kidney disease driven by pathogenic exonic variants [20]. To evaluate whether Volaria’s performance was driven solely by these high-penetrance cases, we assessed the model performance with and without the individuals carrying these variants.

The model accurately predicted outcomes for all individuals carrying monogenic variants, achieving high performance in this small group (e.g., kidney failure AUC = 0.79). Removing carriers of reported monogenic variants had minimal impact on predictive performance of the models in the rest of the patients for all three outcomes, kidney failure, eGFR decline, and steroid resistance. Kidney failure prediction and eGFR decline remained unchanged (ROC AUC = 0.63 for kidney failure and ROC AUC = 0.67 for eGFR decline). Steroid resistance showed a minor increase after excluding monogenic cases (AUC = 0.58) (Supp. Table 1). This demonstrates that Volaria captures both known monogenic disease signals and other novel predictive features, capturing broad polygenic and regulatory contributions to kidney disease progression.

### Volaria is robust across specific disease diagnoses

The CureGN cohort includes individuals diagnosed with one of four rare glomerular diseases (Focal Segmental Glomerulosclerosis (FSGS), IgA Nephropathy (IgAN), Membranous Nephropathy (MN), and Minimal Change Disease (MCD)), each with distinct histopathological characteristics, clinical presentations, and rates of progression, but overlapping disease initiation and progression factors. To understand how predictive performance varies by disease context, we evaluated model performance separately within each diagnosis group for kidney failure and eGFR decline outcomes, the two outcomes relevant to all four diseases. Volaria consistently achieved strong performance for both kidney failure and eGFR decline for FSGS, IgAN, and MCD (though the MCD group’s very small sample size limits interpretability of this result) (Supp. Table 2). Performance in MN was lower, but precision-recall remained comparable for eGFR, suggesting that the model still captures meaningful risk signals and indicating that an MN-specific model would be advantageous as increased sample size becomes available to facilitate its training.

### Cross-cohort generalization

To assess generalizability, we applied Volaria to the GTEx dataset [17] - a large cohort with 440 participants with whole-genome sequencing and clinical annotations. While structured longitudinal outcomes are not available in this cohort, we identified 51 participants with end stage renal disease (annotations for renal failure or dialysis) (Methods). The Volaria model trained on CureGN data achieved renal failure prediction ROC AUC = 0.75, despite differences in cohort design, population and phenotype granularity (Figure 3F). This highlights Volaria’s ability to generalize across cohorts and maintain predictive performance under real-world heterogeneity.

### Feature importance highlights inflammatory and immune pathways

To investigate the biological underpinnings of Volaria’s predictions, we performed SHAP-based [21] feature importance analysis across all outcomes. This approach quantifies the marginal contribution of each feature to model output by estimating changes in predicted score when that feature is perturbed while all others are held constant. We analyzed features with non-zero SHAP values across all random seeds to ensure robustness. For kidney failure prediction, a total of 261 genes consistently contributed to the model output across all diagnosis and genetic backgrounds. To assess whether these features reflect shared molecular mechanisms, we applied Louvain clustering on a tissue-specific functional gene network constructed from a large compendium of functional genomic data [22]. This revealed three functionally coherent modules (Figure 4A). The first module, characterized by IL-1 mediated inflammatory signaling, was enriched for genes involved in IL-1 production and regulation, as well as inositol lipid-mediated signaling, highlighting the contribution of inflammation in progressive kidney injury. The second module represented cellular stress and immune modulation, and included genes linked to cell cycle regulation, DNA damage checkpoints, proteasomal catabolism, and negative regulation of T cell activation, suggesting coordinated stress response and immune regulation mechanisms associated with disease progression. The third module was linked to the negative regulation of signaling receptor activity, and consisted of a smaller set of genes enriched for modulation of receptor signaling processes.

**Figure 4.**
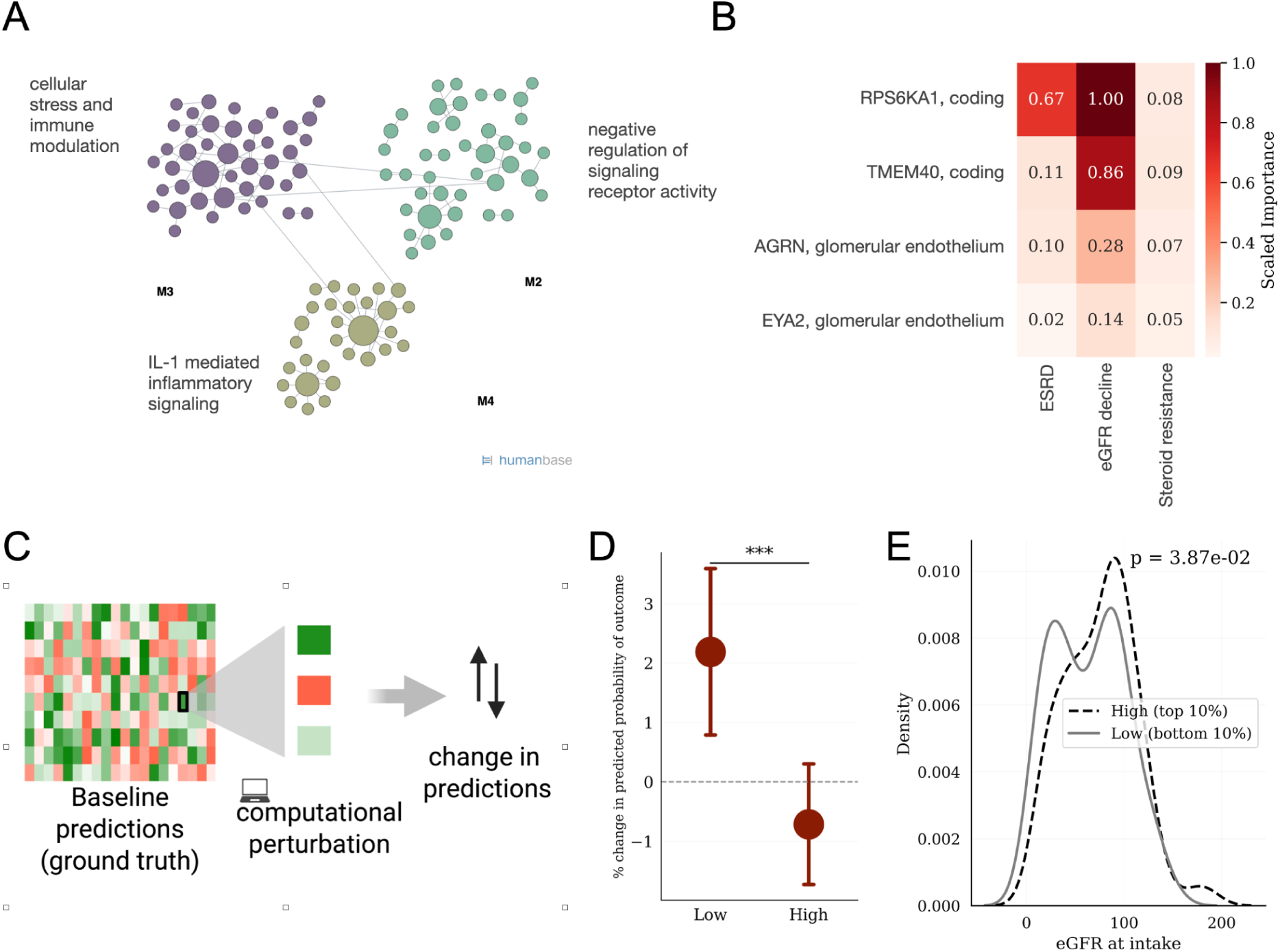
Feature importance and computational perturbation. (A) Functional modules derived from genes with non-zero SHAP-based feature importance for kidney failure prediction, showing enrichment for immune regulation, inflammatory signaling, and modulation of receptor activity. Community detection was performed on SHAP-selected genes using functional networks. (B) Genomic features with non-zero SHAP importance scores in all three outcomes (kidney failure, eGFR decline, and steroid resistance) ranked by scaled median contribution across random seeds. (C) Schematic of the computational perturbation analysis. A single feature is computationally modified within the embedding, and model predictions are recomputed to quantify its isolated contribution (see Methods). (D) Predicted risk of eGFR decline following computational perturbation of the RPS6KA1 coding score. Perturbation analysis indicated a protective effect of strongly deleterious mutations: individuals with less deleterious scores (Low group) showed significantly higher predicted risk than those with highly deleterious values (p = 1.14 × 10⁻²⁴⁰). (E) Measured eGFR at intake for individuals in the cohort within the top and bottom 10% of *RPS6KA1* scores. Lower scores were associated with reduced baseline kidney function (p = 3.87e-02), consistent with the model’s predicted direction of effect.

We next extended the analysis to all three outcomes. Four features were consistently predictive across all three outcomes: coding disruptions in RPS6KA1 (p90 RSK) and TMEM40, and regulatory dysregulations of AGRN and EYA2 in the glomerular endothelium cell type. Specifically, RPS6KA1 showed strong contribution to both kidney failure and eGFR decline, and non-zero contribution to steroid resistance (Figure 4B).

### Computational perturbation reveals protective and adverse gene effects

We then examined whether individual gene-level features can have biologically interpretable effects on clinical outcome prediction by modifying individual embeddings to simulate perturbations affecting specific genes at the regulatory or coding level (Figure 4C). Given its consistently high SHAP contributions, we focused on RPS6KA1 for targeted computational perturbation. We first perturbed the RPS6KA1 coding score, setting it to zero (“Low”) or scaling it up (“High”) across all individuals. The modified embeddings were passed through the fixed outcome model to quantify changes in predicted risk. Individuals with low RPS6KA1 values showed significantly elevated predicted risk of eGFR decline compared to the high group (p = 1.14 × 10⁻²⁴⁰, effect size r = 0.74; Figure 4D), suggesting RPS6KA1 may be protective against eGFR decline. This trend persisted both in training and in the held-out test set (Supp. Figure 3A).

To evaluate concordance of this prediction with clinical data, we stratified cohort participants by their true RPS6KA1 coding scores and compared their measured eGFR at study intake. Individuals in the top decile of RPS6KA1 scores had significantly higher baseline eGFR than those in the bottom decile (p = 0.039; Figure 4E), consistent with the direction of model predictions.

We repeated this analysis in a high-impact regulatory feature (Figure 4B), specifically cell-type specific regulation of EYA2 in the glomerular endothelium. In contrast to the protective effect observed with RPS6KA1, increasing the EYA2 regulatory disruptions in the glomerular endothelium led to significantly higher predicted risk of eGFR decline (Supp. Figure 3 B-C). Consistent with Volaria’s predictions, CureGN participants with high baseline EYA2 scores had lower eGFR at intake (p = 0.05; Supp. Figure 3D). These findings illustrate that Volaria’s structured embeddings encode interpretable, cell type-specific and directionally consistent biological signals.

## Discussion

In this study, we introduced Volaria, a framework for constructing gene-centered representations of genome-wide variation that integrate regulatory and exonic signals into structured, interpretable features. Volaria enables outcome modeling directly from germline whole-genome sequencing, capturing biologically grounded predictive signals without requiring clinical covariates, treatment history, or longitudinal follow-up. While not intended as a replacement for clinical risk models, Volaria provides a complementary layer of genomic insight that can be integrated with phenotypic data in future multi-modal approaches to precision medicine. By encoding both cell type-specific regulatory activity and variant-level pathogenicity into unified representations, Volaria bridges functional variant prediction and genome-based outcome modeling, offering a scalable and reusable foundation for mechanistic hypothesis generation and patient stratification.

Applied to individuals with rare glomerular disease in the CureGN cohort, Volaria robustly predicted treatment resistance and functional decline, outperforming population-level polygenic risk scores and simplified baseline models. Despite the clinical and genetic heterogeneity inherent to glomerular pathophysiology, Volaria maintained performance across diagnosis subgroups, capturing outcome-relevant signals even in underpowered settings. Generalization to the GTEx cohort further confirmed the portability of the learned representations across cohorts, phenotype definitions, and treatment regimens. Although evaluated here in the context of rare glomerular disease, Volaria is a generalizable framework that can be extended to other clinical settings.

Volaria can also be used to identify risk genes, as demonstrated by computational perturbations that revealed interpretable, directional effects of both coding and regulatory features. This capacity to computationally probe the functional landscape of the genome offers a scalable approach for generating testable mechanistic hypotheses and identifying candidate genes for therapeutic targeting.

We note several limitations and potential future directions. Clinical outcomes in longitudinal studies such as CureGN continue to be updated; future follow-up may refine case-control definitions or uncover late-onset events. Treatment, environmental, and social determinants of health, not yet modeled here due to sample size limitations, are also likely to contribute to residual variability. Incorporating these external covariates by applying the Volaria framework to larger datasets could enhance model precision and enable dissection of genome-encoded and external factors influencing clinical outcomes.

While Volaria’s current formulation focuses on six kidney-relevant cell types, the framework is modular and extensible. Additional predictive models spanning cell types, developmental stages, or multi-omic signals could further enhance resolution and biological specificity. Volaria also provides a foundation for genome-to-phenotype modeling in broader settings, including drug response, hospitalization, and rare disease classification. Together, our results establish Volaria as a generalizable and interpretable framework for clinical genome analysis, bridging variant-level sequence predictions with individual-level outcome modeling in a biologically grounded manner.

## Methods

### CureGN cohort

CureGN is a multi-center study with over 1,846 patients with whole-genome sequencing (WGS) and detailed clinical follow-up spanning years. CureGN participants were enrolled and tracked annually at treatment centers with standardized data collection. Participants had kidney tissue biopsy derived disease status and, following enrollment, participants provided blood and urine biosamples. Throughout the course of the study, participants have semi-annual follow-up visits. Key negative endpoints in the program are kidney function loss, defined as a 40% estimated glomerular filtration rate (eGFR) decline and kidney failure (defined in the project as having either two consecutive eGFR <15, kidney transplant, or chronic dialysis). Individuals can also be labeled as steroid resistant based on their proteinuria response to steroid treatment, representing a non-temporal outcome.

### Variant-level information

ExpectoSC scores were computed for all the variants in the range of +/− 20kbp of the gene TSS. The location of the TSS was selected based on the top CAGE [13,23] peak if present, or else GENCODE TSS [24] (Release 19, GRCh37.p13), in agreement with the ExpectoSC pipeline [14]. Coding variants were defined as SNVs within annotated exons in GENCODE. AlphaMissense VEP [25] plugin was used to get AlphaMissense scores [15]. The variant file with patient genotype information was obtained from the CureGN consortium. The genotypes with read depth (DP) less than 10 or genotype quality (GQ) under 20 were filtered out. In addition, Variant Quality Score Recalibration (VQSR) from the Genome Analysis Toolkit (GATK) [26] was used to filter variants to the 99.8 sensitivity tranche for SNVs. No further AF/MAF filtering was performed. All annotations were harmonized to GRCh37.

### Combining variant effects at the gene level

Predictions of each variant were computed independently, and then combined per gene by summing the scores and normalizing by the number of variants in the gene:

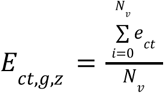

where *N*_*v*_ is the total number of variants in gene *g*, *e*_*ct*_ is the gene dysregulation effect for the cell type *ct* and *z* is the zygosity. The effects for the exonic variants (also referred to as coding throughout the text), were not cell type specific and represented by AlphaMissense score. Homozygous and heterozygous variants were processed separately first, and then aggregated as *E*_*ct*,*g*_ = |*E*_*ct*,*g*,*homoz*_ + 0. 5*E*_*ct*,*g*,*heteroz*_|.

### Individual matrix representation

After combining the predictions, each patient *p*_*i*_ had two primary sets of representations: regulatory set and variants located in the exonic regions. Regulatory effects were further separated into cell types. The final gene modulation matrix *P*_*G*_ was *N* ×*M*, where *N* is the number of cell types plus exonic variant predictors, *M* is the number of genes, and *i^th^* row of the matrix represents *i^th^* cell type:

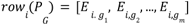

### Application to GTEx and CureGN

To identify individuals with kidney failure in the GTEx cohort, we used the labels MHDLYSIS (dialysis) and MHRNLFLR (renal failure). Additional labels including MNNEPH (nephritis), MHUREMIA (uremia), and MHGENCMT (clinical notes with kidney-related terms) were used to define a kidney-impacted subset. The remaining individuals (n=378) served as the background set. ExpectoSC kidney models for podocyte, myofibroblast, glomerular endothelium, B cell, CD4 T cell, CD8 T cell were used. In the CureGN cohort, outcomes were represented as binary labels; kidney failure and eGFR also had a corresponding “_time” feature representing event or follow-up time. To ensure rigorous separation of training and test data, the test set (20%) was created via stratified sampling into five bins based on kidney failure time and held out from all subsequent modeling decisions. All feature selection steps, including collinearity removal and filtering, were performed exclusively on training data. Hyperparameter optimization via cross-validation was conducted entirely within the training set. SHAP analysis was performed post-hoc for interpretation and did not influence feature selection or model architecture. To account for population structure, the first 10 principal components (PCs) were regressed out separately for each feature matrix *E*_*i*_ for the individuals in the background GTEx cohort. Features with no predicted impact in any of the individuals in the CureGN train set were dropped and data was centered using z-score normalization: 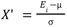, followed by computing top 10 eigenvectors to get a projection matrix *P*_*i*_ = *X*’*W*, where *W* is the eigenvectors of the covariance matrix 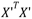. Then a linear regression model was fit, such that *Y*_*i*, *pred*_ = *P*_*i*_β_*i*_ + ɛ_*i*_, where β are the regression coefficients. Finally, the residuals were computed as *Y*_*i*, *res*_ = |*Y*_*i*_ − *Y*_*i*, *pred*_|. All steps were implemented using `scikit-learn` [27]. The transformations were fitted on the background GTEx set. Initial features selection was done by first dropping collinear features (threshold=0.9) using train split, and features with median scores above 99th percentile were retained. *Fitting models for outcome prediction*: RandomForestClassifier with balanced class weights and entropy criterion was used, using `scikit-learn` implementation. The number of estimators and maximum depth was found using 5-fold 5 repeat cross validation across, *max*_*depth* = [2, 5, 7]; *n*_*estim* = [100, 500, 1000], separately for each of the three outcomes. In test set, individuals with less than 2000 days of follow up and no event registered were dropped, and decision to drop them from training was made with cross validation: *filter* = [*True*, *False*].

The resulting best parameters were: *kidney failure*: [*True*, 2, 1000]; *eGFR*_*drop*: [*True*, 2, 100]; *Steroid resistance*: [*True*, 2, 1000]. After cross-validation, the models were trained on the train set with 7 different random states. ROC AUC and AP (average_precision_score) was computed with the `scikit-learn, and` gain over baseline in AP was computed as (*s*_*model*_ − *s*_*baseline*_)/*s*_*baseline*_.

### Evaluating embedding structure contribution

To benchmark against Volaria’s structured, gene-and cell type–resolved embeddings, we constructed flat baselines by collapsing each feature set to per-individual scalars after the same feature selection used for the main models. For the coding baseline, we took the maximum absolute coding-effect value across all genes. For the regulatory baseline, we took the maximum absolute regulatory-effect value across all genes and cell types. We evaluated four input settings: sex only; coding scalar only; regulatory scalar only; and a two-scalar combination of coding + regulatory. Inputs were standardized on the training split and applied to the test split. Models matched the main setup, re-fit across multiple random seeds and evaluated on the same fixed test split using ROC AUC and AP gain; results are reported as mean ± standard deviation across seeds.

### PGS computation

We used pgs_calc package [28] with nextflow [29]. Public PGS scoring files were retrieved from the PGS Catalog using the relevant PGS IDs. For each individual, scores were computed with the weights provided by the catalog, without modification. All PGS were derived under the GRCh37 build. The resulting PGS were then input into the same downstream prediction models used for Volaria, with identical parameters and train / test splits.

### Feature importance

We computed SHAP values to estimate feature contributions to model predictions using the TreeExplainer from the SHAP library [21]. The explainer was initialized with the trained model and a background sample consisting of the training set. We used the interventional feature perturbation approach and set the model output to probability. Feature importance was calculated as the mean absolute SHAP value across samples for the positive class, where each feature is a combination of the gene affected and context. Importances were computed separately for each model across random seeds and outputs. Features that had non-zero weights in all random seeds were selected.

### Computational perturbation

Computational perturbation was performed by modifying the embedding of a given feature within its context. For the Low setting, the embedding was set to zero; for the High setting, it was scaled by a factor of 100 relative to the ground truth embedding. Predictions from the modified embeddings were normalized using the formula: 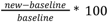. To assess phenotypic impact, individuals were stratified into top and bottom 10% based on the ground truth embedding using np.quantile, and those without eGFR measurements were excluded afterward, resulting in 80 individuals in the bottom group and 83 in the top group. Group distributions were compared using the Mann–Whitney U test, and effect size was calculated as Rosenthal’s 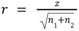, where *z* is standardized U value under the null hypothesis and *n*_1_ and *n*_2_ are the sizes of the respective groups.

## Data Availability

GTEx whole genome data is available through dbGaP (phs000424.v10.p2); CureGN data is available by request through the CureGN consortium. De-identified data generated in this study may be available for research purposes from the corresponding authors upon reasonable request and is subject to institutional and consortium data use agreements.

## Code Availability

Code to run Volaria and reproduce all figures is available on GitHub https://github.com/ksenia007/volaria

## Contributions

K.S. and O.G.T. conceived the study. K.S. designed the study, implemented the framework, performed analyses, and wrote the manuscript. O.G.T. supervised the project, contributed to its conceptual development and manuscript. L.M. and M.K. contributed clinical expertise and guidance on disease context and study interpretation. V.N.K. contributed to conceptual discussions, manuscript development, and study direction. C.T. provided input on biological interpretation. C.Y.P contributed to the conceptual discussions and manuscript development. All authors reviewed and approved the final manuscript.

## Acknowledgements

This work was supported in part by the National Institutes of Health (NIH) under grant R01GM071966 to O.G.T., and by subawards from NIH U01DK133090 and NIH RC2DK116690 to O.G.T.

We gratefully acknowledge the CureGN Consortium for access to the genomic and clinical data used in this study, as well as for continued collaboration and feedback. We specifically thank Krzysztof Kiryluk and Chen Wang for their work on variant calling and data processing. Funding for the CureGN consortium is provided by U24DK100845, U01DK100846, U01DK100876, U01DK100866, and U01DK100867 from the National Institute of Diabetes and Digestive and Kidney Diseases (NIDDK). Patient recruitment is supported by NephCure. Dates of funding for first phase of CureGN was 9/16/2013-5/31/2019. Dates of funding for the second phase of CureGN was 6/1/2019 - 5/31/2024.

We thank members of the Troyanskaya lab for their helpful feedback and discussions during the preparation of this manuscript.

A substantial portion of this research was performed using the Princeton Research Computing resources at Princeton University, a consortium led by the Princeton Institute for Computational Science and Engineering (PICSciE) and the Office of Information Technology’s Research Computing group.

Figures 1 and parts of Figures 2 and 4 were generated using BioRender.com

## Supplement

**Supp. Table 1.**
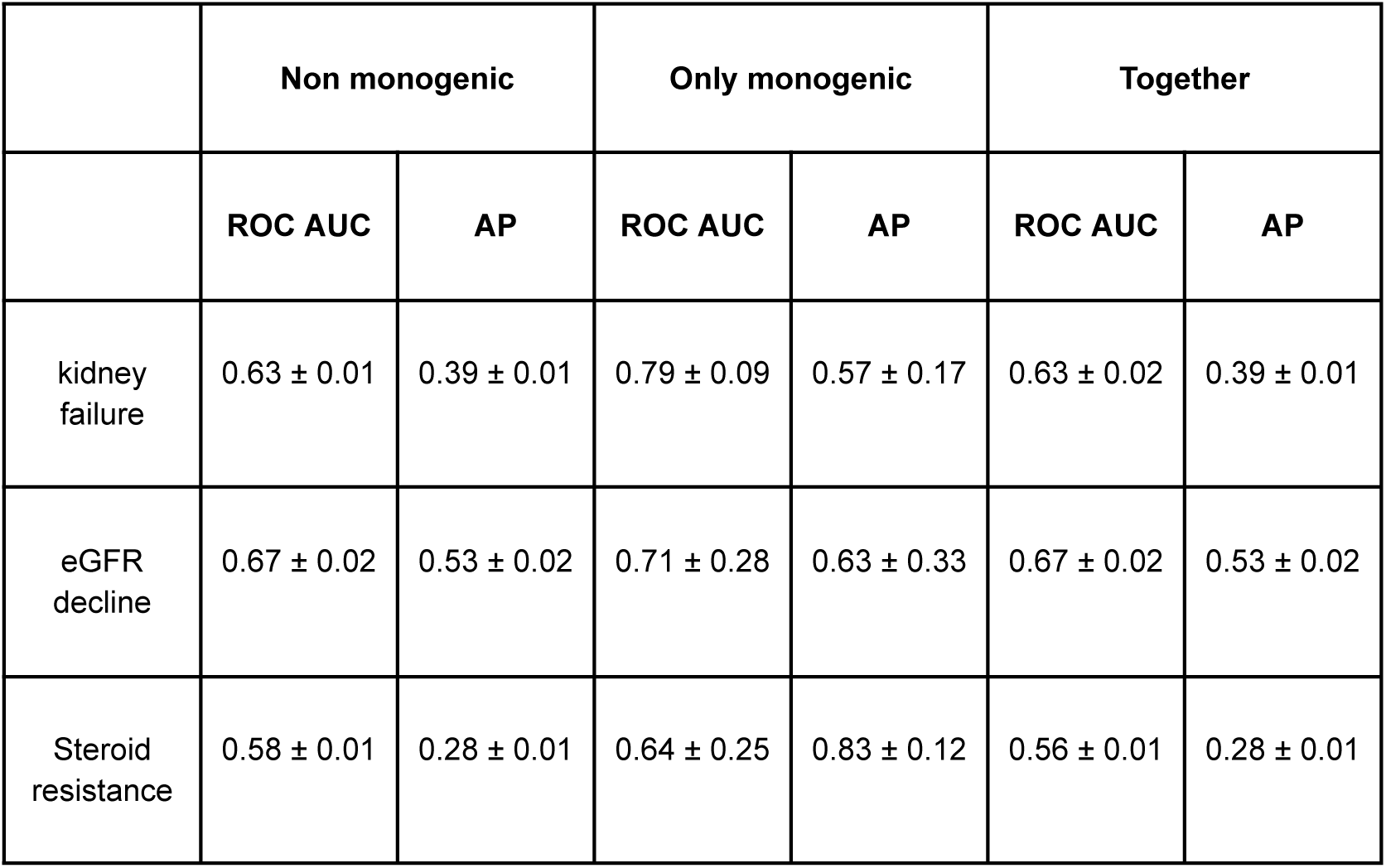
Model performance on individuals with monogenic kidney disease and after their exclusion. Metrics shown: ROC AUC and AP for each outcome in the full test set, non-monogenic subset, and monogenic-only subset. Error is one standard deviation across random seeds.

**Supp. Figure 1.**
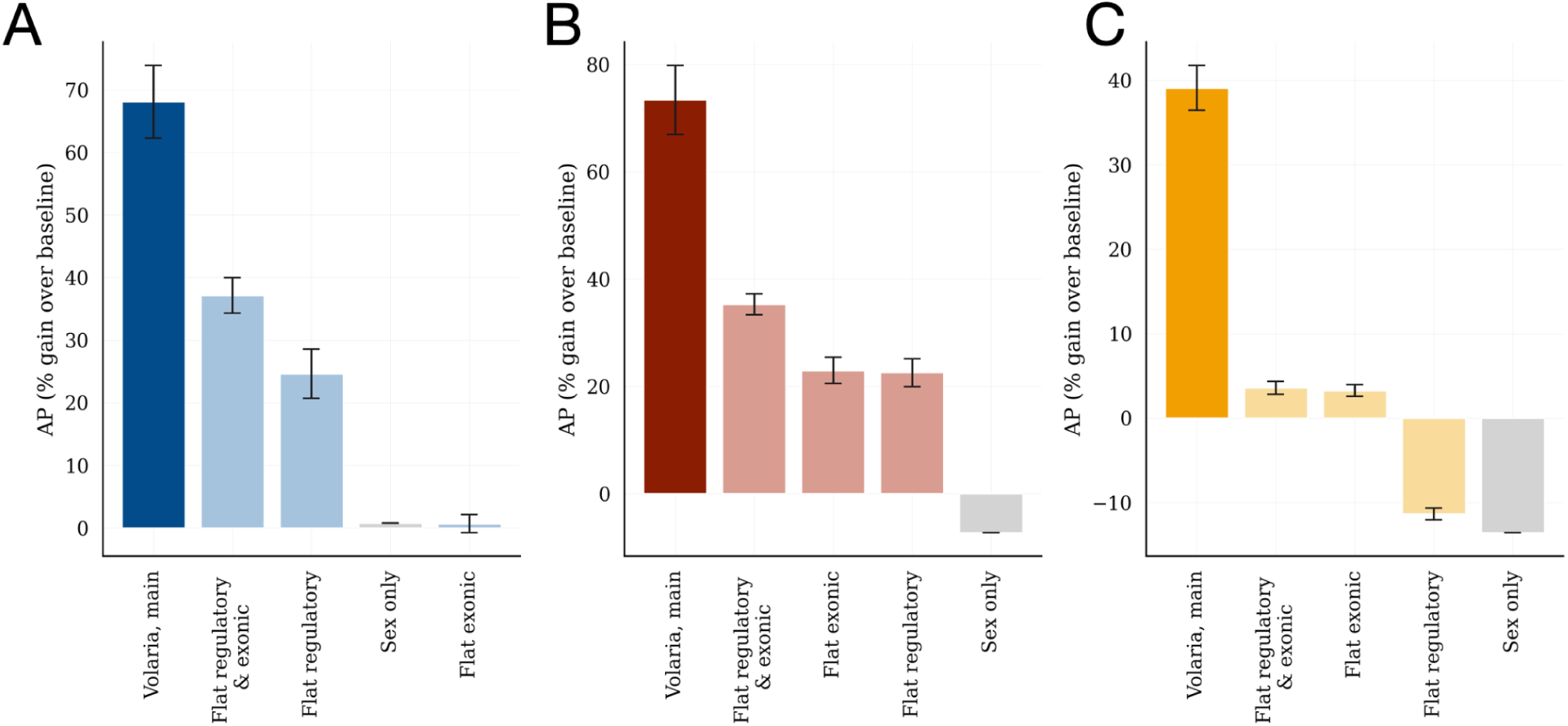
Average precision gains across baseline models. (A–C) Volaria shows improved precision-recall performance over baseline models across three renal outcomes: (A) kidney failure, (B) 40% eGFR decline, and (C) steroid resistance. Values reflect the gain in average precision (AP) over a naive baseline equal to the prevalence of positive cases. Baselines include flat aggregations of functional scores across matched feature sets, separately for regulatory and exonic components, as well as combined, and a clinical-only model using sex.

**Supp. Figure 2.**
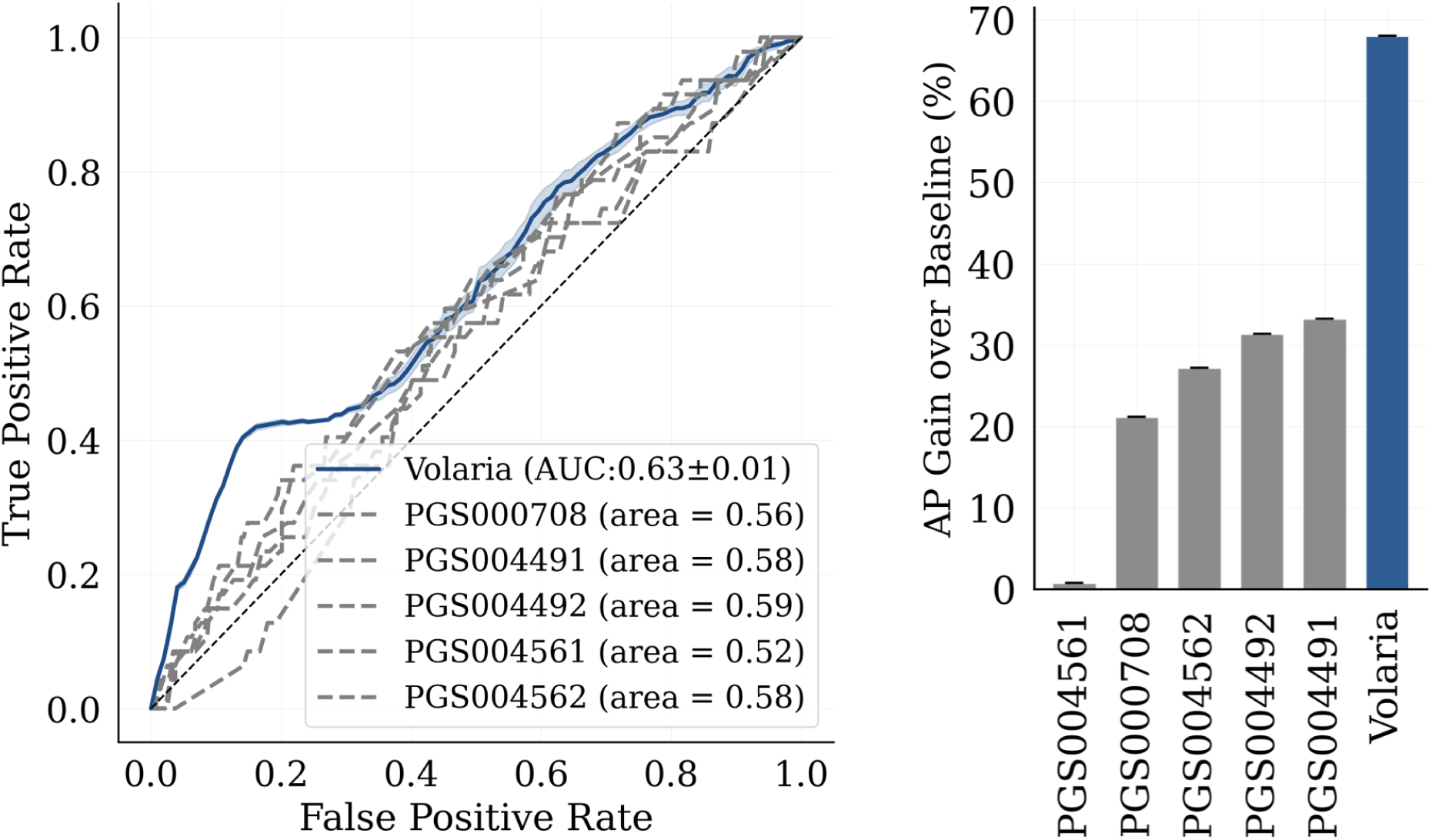
(left) ROC AUC curves for predicting kidney failure outcome using five polygenic scores (PGS) from the PGS Catalog; (right) Average precision improvements over baseline for the kidney failure using each of the PGS scores. Volaria achieves over twice the AP of the best-performing PGS

**Supp. Table 2.** Model performance by diagnosis subgroup for each predicted clinical outcome. Reported metrics are total number of events in the test set, event rates in test and training sets, ROC AUC, AP, and percent gain in AP over baseline. Subgroup-specific metrics highlight variation in outcome prevalence and performance, with high scores observed across both common and rare subtypes.

| Predicted feature | Group | Total events in test | Event rate in test (%) | Event rate in train (%) | ROC AUC | AP | Gain in AP (%) |
| --- | --- | --- | --- | --- | --- | --- | --- |
| eGFR decline | ALL | 63 | 30.6 | 30.9 | 0.67 ± 0.02 | 0.53 ± 0.02 | 73.4 ± 6.4 |
| eGFR decline | FSGS | 22 | 40.0 | 47.6 | 0.69 ± 0.03 | 0.64 ± 0.02 | 59.9 ± 6.0 |
| eGFR decline | IgAN | 20 | 28.6 | 23.7 | 0.69 ± 0.04 | 0.59 ± 0.03 | 105.0 ± 11.8 |
| eGFR decline | MN | 17 | 38.6 | 37.4 | 0.59 ± 0.05 | 0.58 ± 0.02 | 49.4 ± 4.9 |
| eGFR decline | MCD | 4 | 10.8 | 16.4 | 0.74 ± 0.06 | 0.39 ± 0.07 | 263.8 ± 61.4 |
| kidney failure | ALL | 47 | 23.0 | 26.1 | 0.63 ± 0.02 | 0.39 ± 0.01 | 68.1 ± 5.8 |
| kidney failure | IgAN | 18 | 25.7 | 21.3 | 0.68 ± 0.02 | 0.56 ± 0.01 | 118.3 ± 5.0 |
| kidney failure | FSGS | 17 | 30.9 | 47.9 | 0.68 ± 0.02 | 0.48 ± 0.02 | 56.9 ± 4.9 |
| kidney failure | MN | 10 | 23.8 | 21.6 | 0.46 ± 0.04 | 0.38 ± 0.01 | 60.0 ± 3.9 |
| kidney failure | MCD | 2 | 5.4 | 11.6 | 0.73 ± 0.04 | 0.18 ± 0.02 | 241.6 ± 35.5 |
| Steroid resistance | ALL | 40 | 19.9 | 19.2 | 0.56 ± 0.01 | 0.28 ± 0.01 | 39.1 ± 2.6 |
| Steroid resistance | FSGS | 21 | 39.6 | 33.2 | 0.42 ± 0.03 | 0.39 ± 0.01 | -2.5 ± 3.3 |
| Steroid resistance | MCD | 12 | 28.6 | 29.0 | 0.68 ± 0.02 | 0.48 ± 0.02 | 69.4 ± 8.5 |
| Steroid resistance | IgAN | 4 | 6.2 | 7.5 | 0.70 ± 0.04 | 0.27 ± 0.07 | 339.4 ± 107.1 |
| Steroid resistance | MN | 3 | 7.3 | 11.7 | 0.77 ± 0.05 | 0.24 ± 0.05 | 230.0 ± 63.0 |

**Supp. Figure 3.**
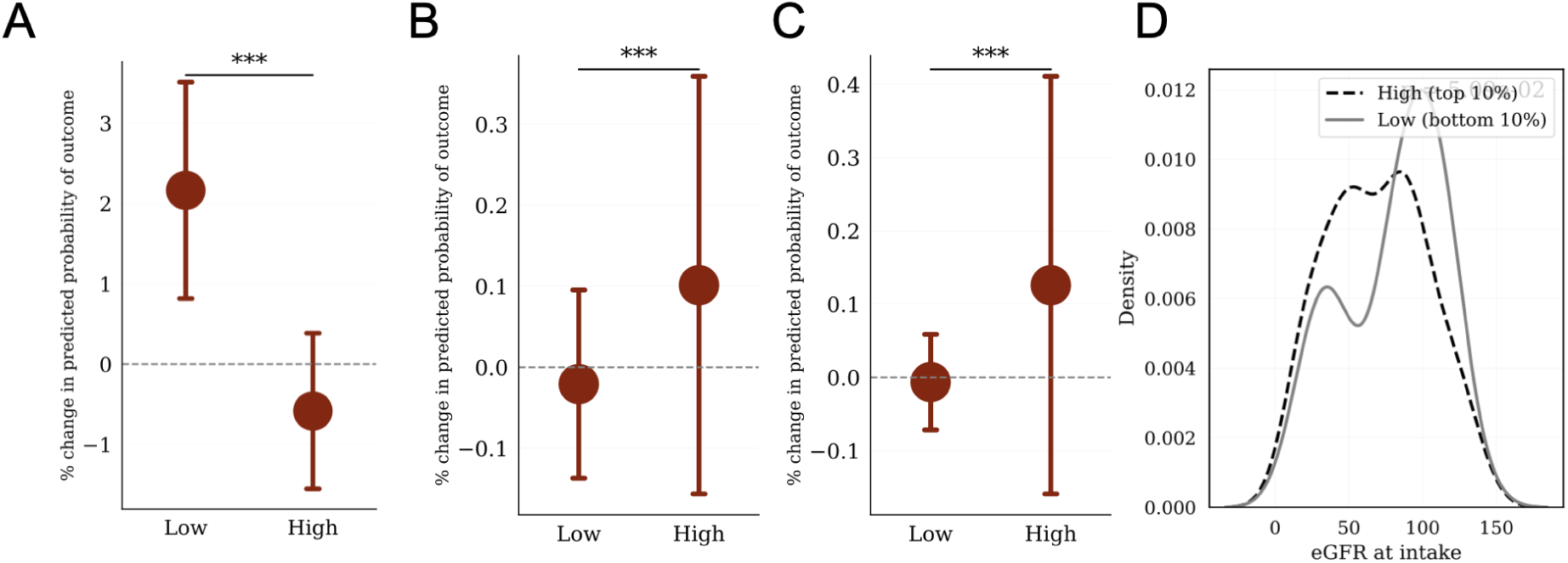
Computational perturbation of the embeddings. (A) Perturbation of the RPS6KA1 coding score in the test set showed significant differences in predicted eGFR decline risk between high and low conditions, in agreement with train subset results. (B) In silico modification of the EYA2 regulatory score in the glomerular endothelium, training set subset. Higher values led to increased predicted risk of eGFR decline. (C) EYA2 computational perturbation in the glomerular endothelium repeated in the test set showed consistent results, with higher scores again associated with worse predictions. (D) Individuals with high baseline non-modified EYA2 regulatory scores had lower eGFR at intake, supporting the adverse directionality observed in the computational perturbations.

## CureGN Collaborators

The CureGN Consortium members listed below, from within the four Participating Clinical Center networks and Data Coordinating Center, are acknowledged by the authors as Collaborators.

**CureGN Principal Investigators; *CureGN Site Principal Investigators; ^+^CureGN Pathologists, ^#^CureGN Lead Coordinators.

### CureGN Participating Clinical Centers (PCC) through Columbia University

*<u>Columbia University, New York, NY, US:</u>* Gerald Appel, Revekka Babayev, Ibrahim Batal ^+^, Andrew Bomback**, Pietro Canetta, Brenda Chan, Vivette Denise D’Agati ^+^, Samitri Dogra, Hilda Fernandez, Gabriele Gaggero^+^, Ali Gharavi**, William Hines, Syed Ali Husain, Krzysztof Kiryluk**, Satoru Kudose ^+^, Fangming Lin, Victoria Kolupaeva#, Maddalena Marasa, Glen Markowitz ^+^, Mariela Naarro-Torres, Hila Milo Rasouly, Sumit Mohan, Nicola Mongera, Jordan Nestor, Thomas Nickolas, Jai Radhakrishnan, Maya Rao, Maya Sabatello, Simone Sanna-Cherchi, Dominick Santoriello^+^, Miroslav Sekulic ^+^, Shayan Shirazian, Michael Barry Stokes^+^, Natalie Uy, Natalie Vena, Benjamin Wooden

*<u>University of Warsaw, Warszawa, Poland:</u>* Bartosz Foroncewicz, Natalia Krata, Barbara Moszczuk, Krzysztof Mucha*, Agnieszka Perkowska-Ptasińska, Elżbieta Ryszkowska

*<u>IRCCS Giannina Gaslini, Genoa, Italy:</u>* Gian Marco Ghiggeri*, Francesca Lugani, Valerio Vellone^+^

### CureGN Participating Clinical Centers (PCC) through the Pediatric Nephrology Research Consortium

*<u>Children’s Hospital of Michigan, Detroit, MI, USA:</u>* Rossana Baracco, Amrish Jain*

*<u>Children’s Hospital of New Orleans/ LSU Health, New Orleans, LA, USA:</u>* Diego Aviles*

*<u>Children’s Mercy Hospital, Kansas City, MO, USA:</u>* Tarak Srivastava*, Alexander Katz^+^

*<u>Children’s National Medical Center, Wmcgashington DC, USA:</u>* Sun-Young Ahn*

*<u>Cincinnati Children’s Hospital Cincinnati, OH, USA</u>*: Prasad Devarajan, Elif Erkan*, Donna Claes, Hillarey Stone

*<u>Connecticut Children’s Medical Center, Hartford, CT, USA:</u>* Sherene Mason*

*<u>East Carolina University Brody School of Medicine, Greenville, NC, USA:</u>* Liliana Gomez-Mendez*

*<u>Emory University, Atlanta, GA, USA:</u>* Larry Greenbaum**, Chia-shi Wang, Hong (Julie) Yin^+^

*<u>Helen DeVos Children’s Hospital, Grand Rapids, MI, USA:</u>* Goebel Jens*, Julia_Steinke

*<u>Levine Children’s Hospital/Atrium Health, Charlotte, NC, USA:</u>* Donald Weaver*

*<u>Lurie Children’s Hospital, Chicago IL, USA:</u>* Jerome Lane*

*<u>Mayo Clinic, Rochester, MN, USA:</u>* Carl Cramer*

*<u>Medical College of Wisconsin, Milwaukee, WI, USA:</u>* Cindy Pan, Neil Paloian, Rajasree Sreedharan*

*<u>Medical University of South Carolina, Charleston SC, USA:</u>* David Selewski, Katherine Twombley*, Sally Self^+^

*<u>Nationwide Children’s Hospital, Columbus, OH, USA:</u>* Samantha Martinek-Bundt#, Dawson Carmean#, Mary Dreher^#^, Aria Dockham^#^, Mahmoud Kallash*, John Mahan, Samantha Sharpe^#^, William Smoyer**, Laura Biederman^+^

*<u>Oregon Health and Science University, Portland, OR, USA:</u>* Amira Al-Uzri*, Sandra Iragorri

*<u>Riley Children’s Hospital, Indianapolis, IN, USA:</u>* Myda Khalid**

*<u>Cardinal Glennon Children’s Medical Center/ St. Louis University, St. Louis, MO, USA:</u>* Craig Belsha*

*<u>Texas Children’s Hospital, Houston, TX, USA</u>*: Elizabeth Onugha*, Michael Braun, AC Gomez

*<u>Texas Tech Health Sciences Center, Amarillo, TX, USA:</u>* Tetyana Vasylyeva*

*<u>Children’s of Alabama, University of Alabama, Birmingham, AL, USA:</u>* Daniel Feig*

*<u>University of Colorado Children’s Hospital, Colorado, Aurora, CO, USA:</u>* Melisha Hannah*

*<u>University of Iowa Children’s Hospital, Iowa City, IA, USA:</u>* Carla Nester*

*<u>University of Kentucky, Lexington, KY, USA:</u>* Aftab Chishti*

*<u>University of Louisville, Louisville, KY, USA:</u>* Jon Klein**

*<u>Holtz Medical Center, University of Miami, Miami, FL, USA:</u>* Chryso Katsoufis, Wacharee Seeherunvong*

*<u>University of Minnesota Children’s Hospital, Minneapolis, MN, USA:</u>* Michelle Rheault**

*<u>University of New Mexico Health Sciences Center, Albuquerque, NM, USA:</u>* Craig Wong*

*<u>University of Oklahoma Health Sciences Center, Oklahoma City, OK, USA:</u>* Qassim Abid*

*<u>University of Virginia, Charlottesville, VA, USA:</u>* John Barcia*, Agnes Swiatecka-Urban

*<u>University of Wisconsin, Madison, WI, USA:</u>* Sharon Bartosh*

*<u>Vanderbilt Children’s Hospital, Nashville TN, USA:</u>* Tracy Hunley*

*<u>Washington University in St. Louis, St. Louis, MO, USA:</u>* Vikas Dharnidharka*, Brian Stotter, Joseph, Gaut ^+^

### CureGN Participating Clinical Centers (PCC) through the University of North Carolina

*<u>Hôpital Maisonneuve-Rosemont, Montreal, Canada:</u>* Louis-Philippe Laurin*, Virginie Royal^+^, Mathieu Latour^+^, Natlie (Natacha) Patey ^+^

*<u>Medical University of South Carolina, Charleston, SC, USA:</u>* Anand Achanti, Milos Budisavljevic*

*<u>Northwestern University, Chicago, IL, USA:</u>* Cybele Ghossein, Yonatan Peleg *

*<u>Ohio State University, Columbus, OH, USA:</u>* Salem Almaani*, Isabelle Ayoub, Samir Parikh, Brad Rovin, Anjali Satoskar^+^

*<u>University of Chicago, Chicago, IL, USA:</u>* Anthony Chang^+^

*<u>University of Alabama at Birmingham, Birmingham, AL, USA:</u>* Huma Fatima^+^, Jan Novak, Matthew Renfrow, Dana Rizk*

*<u>University of North Carolina Kidney Center, Chapel Hill, NC, USA</u>*: Dhruti Chen, Vimal Derebail**, Ronald Falk**, Keisha Gibson, Dorey Glenn, Susan Hogan, Koyal Jain, J. Charles Jennette^+^, Amy Mottl, Caroline Poulton^#^, Monica Reynolds, Manish Kanti Saha, Nicole E. Wyatt

*<u>Vanderbilt University, Nashville, TN, USA:</u>* Agnes Fogo^+^, Neil Sanghani*

*<u>Virginia Commonwealth University, Richmond, VA, USA:</u>* Jason Kidd*, Selvaraj Muthusamy^+^

### CureGN Participating Clinical Centers (PCC) through the University of Pennsylvania

*<u>Children’s Hospital of Philadelphia, Philadelphia, PA, USA</u>*: Michelle Denburg, Amy Kogon, Kevin Meyers*, Madhura Pradhan

*<u>Cleveland Clinic, Cleveland, OH, CA:</u>* Raed Bou Matar*, John O’Toole, John Sedor

*<u>Cohen Children’s Medical Center, New Hyde Park, NY, USA:</u>* Christine Sethna*^, Suzanne Vento^#^

*<u>Johns Hopkins University, Baltimore, MD, USA:</u>* Mohamed Atta, Serena Bagnasco^+^, Alicia Neu, John Sperati*

*<u>Lundquist Institute at Harbor-UCLA Medical Center, Torrance, CA, USA:</u>* Sharon Adler*, Tiane Dai, Ram Dukkipati

*<u>Mayo Clinic, Rochester, MN, USA:</u>* Fernando Fervenza*, Sanjeev Sethi ^+^

*<u>Montefiore Medical Center, The Bronx, New York, NY, USA:</u>* Frederick Kaskel, Kaye Brathwaite, Kimberly Reidy*

*<u>New York University, New York, NY, USA</u>*: Joseph Weisstuch, Ming Wu ^+^, Olga Zhdanova

*<u>Spokane Providence Medical Center, Spokane, WA, USA:</u>* Katherine Tuttle*

*<u>Stanford University, Palo Alto, CA, USA</u>*: Jill Krissberg, Richard Lafayette*, Kamal Fahmeedah, Elizabeth Talley

*<u>Sunnybrook Health Sciences Centre, Toronto, Canada:</u>* Michelle Hladunewich*

*<u>The Hospital for Sick Children, Toronto, Canada:</u>* Rulan Parekh*

*<u>University Health Network, Toronto, Canada</u>*: Carmen Avila-Casado^+^, Daniel Cattran*, Reich Heather, Philip Boll

*<u>University of Miami, Miami, FL, USA:</u>* Yelena Drexler, Alessia Fornoni*

*<u>University of Michigan, Ann Arbor, MI, USA:</u>* Jeffrey Hodgin^+^, Andrea Oliverio*

*<u>University of Pennsylvania, Philadelphia, PA, USA</u>*: Jon Hogan, Lawrence Holzman**, Matthew Palmer ^+^, Gaia Coppock

*<u>University of Pittsburgh School of Medicine, Pittsburgh, PA, USA:</u>* Michael Mortiz*

*<u>University of Washington, Seattle, WA, USA:</u>* Charles Alpers^+^, J. Ashley Jefferson*

*<u>UT Southwestern, Dallas, TX, USA</u>*: Kamal Sambandam*, Bethany Roehm

### Data Coordinating Center (DCC)

*<u>Cedar Sinai Medical Center, Los Angeles, CA, USA:</u>* Cynthia Nast^+^, Jean Hou^+^

*<u>Duke University, Durham, NC, USA:</u>* Laura Barisoni

*<u>Cleveland Clinic, Cleveland, OH, USA:</u>* Crystal Gadegbeku**

*<u>Northwestern University, Chicago, IL, USA:</u>* Abigail Smith**

*<u>University of Michigan, Ann Arbor, MI, USA:</u>* Brenda Gillespie, Bruce Robinson, Matthias Kretzler, Zubin Modi, Laura Mariani**

### Steering Committee Chair

Lisa M. Guay-Woodford, Children’s Hospital of Pennsylvania, Philadelphia, PA, USA

## References

1. Whole-genome sequencing of 490,640 UK Biobank participants. Nature. 2025; 1–10.

2. The 100, 000 Genomes Project Pilot Investigators. 100,000 Genomes Pilot on Rare-Disease Diagnosis in Health Care — Preliminary Report. New England Journal of Medicine. 2021 [cited 7 Aug 2025]. doi:10.1056/NEJMoa2035790

3. Sherman CA, Claw KG, Lee S-B. Pharmacogenetic analysis of structural variation in the 1000 genomes project using whole genome sequences. Scientific Reports. 2024;14: 1–13.

4. Leong IUS, Cabrera CP, Cipriani V, Ross PJ, Turner RM, Stuckey A, et al. Large-Scale Pharmacogenomics Analysis of Patients With Cancer Within the 100,000 Genomes Project Combining Whole-Genome Sequencing and Medical Records to Inform Clinical Practice. Journal of Clinical Oncology. 2025 [cited 7 Aug 2025]. doi:10.1200/JCO.23.02761

5. Wray NR, Goddard ME, Visscher PM. Prediction of individual genetic risk to disease from genome-wide association studies. Genome Res. 2007;17: 1520–1528.

6. Lek M, Karczewski KJ, Minikel EV, Samocha KE, Banks E, Fennell T, et al. Analysis of protein-coding genetic variation in 60,706 humans. Nature. 2016;536: 285–291.

7. Karczewski KJ, Francioli LC, Tiao G, Cummings BB, Alföldi J, Wang Q, et al. The mutational constraint spectrum quantified from variation in 141,456 humans. Nature. 2020;581: 434–443.

8. Boyle EA, Li YI, Pritchard JK. An expanded view of complex traits: from polygenic to omnigenic. Cell. 2017;169: 1177.

9. Dornbos P, Koesterer R, Ruttenburg A, Nguyen T, Cole JB, Leong A, et al. A combined polygenic score of 21,293 rare and 22 common variants improves diabetes diagnosis based on hemoglobin A1C levels. Nature Genetics. 2022;54: 1609–1614.

10. Biological insights from 108 schizophrenia-associated genetic loci. Nature. 2014;511: 421–427.

11. Fritsche LG, Igl W, Bailey JN, Grassmann F, Sengupta S, Bragg-Gresham JL, et al. A large genome-wide association study of age-related macular degeneration highlights contributions of rare and common variants. Nature genetics. 2016;48. doi:10.1038/ng.3448

12. Chen KM, Wong AK, Troyanskaya OG, Zhou J. A sequence-based global map of regulatory activity for deciphering human genetics. Nat Genet. 2022;54: 940–949.

13. Avsec Ž, Agarwal V, Visentin D, Ledsam JR, Grabska-Barwinska A, Taylor KR, et al. Effective gene expression prediction from sequence by integrating long-range interactions. Nature Methods. 2021;18: 1196–1203.

14. Sokolova K, Theesfeld CL, Wong AK, Zhang Z, Dolinski K, Troyanskaya OG. Atlas of primary cell-type-specific sequence models of gene expression and variant effects. Cell Rep Methods. 2023;3: 100580.

15. Cheng J, Novati G, Pan J, Bycroft C, Žemgulytė A, Applebaum T, et al. Accurate proteome-wide missense variant effect prediction with AlphaMissense. Science. 2023 [cited 19 Mar 2025]. doi:10.1126/science.adg7492

16. Mariani LH, Bomback AS, Canetta PA, Flessner MF, Helmuth M, Hladunewich MA, et al. CureGN Study Rationale, Design, and Methods: Establishing a Large Prospective Observational Study of Glomerular Disease. American journal of kidney diseases : the official journal of the National Kidney Foundation. 2018;73: 218.

17. Lonsdale J, Thomas J, Salvatore M, Phillips R, Lo E, Shad S, et al. The Genotype-Tissue Expression (GTEx) project. Nature Genetics. 2013;45: 580–585.

18. Sinnott-Armstrong N, Tanigawa Y, Amar D, Mars N, Benner C, Aguirre M, et al. Genetics of 35 blood and urine biomarkers in the UK Biobank. Nature Genetics. 2021;53: 185–194.

19. Jung H, Jung H-U, Baek EJ, Kwon SY, Kang J-O, Lim JE, et al. Integration of risk factor polygenic risk score with disease polygenic risk score for disease prediction. Communications Biology. 2024;7: 1–13.

20. Elliott MD, Vena N, Marasa M, Cocchi E, Bheda S, Bogyo K, et al. Increased risk of kidney failure in patients with genetic kidney disorders. The Journal of Clinical Investigation. 2024;134: e178573.

21. Lundberg SM, Erion G, Chen H, DeGrave A, Prutkin JM, Nair B, et al. From local explanations to global understanding with explainable AI for trees. Nature Machine Intelligence. 2020;2: 56–67.

22. Krishnan A, Zhang R, Yao V, Theesfeld CL, Wong AK, Tadych A, et al. Genome-wide prediction and functional characterization of the genetic basis of autism spectrum disorder. Nature Neuroscience. 2016;19: 1454–1462.

23. Bertin N, Mendez M, Hasegawa A, Lizio M, Abugessaisa I, Severin J, et al. Linking FANTOM5 CAGE peaks to annotations with CAGEscan. Scientific Data. 2017;4: 1–8.

24. Harrow J, Frankish A, Gonzalez JM, Tapanari E, Diekhans M, Kokocinski F, et al. GENCODE: the reference human genome annotation for The ENCODE Project. Genome Res. 2012;22. doi:10.1101/gr.135350.111

25. McLaren W, Gil L, Hunt SE, Riat HS, Ritchie GRS, Thormann A, et al. The Ensembl Variant Effect Predictor. Genome Biol. 2016;17: 1–14.

26. McKenna A, Hanna M, Banks E, Sivachenko A, Cibulskis K, Kernytsky A, et al. The Genome Analysis Toolkit: a MapReduce framework for analyzing next-generation DNA sequencing data. Genome Res. 2010;20. doi:10.1101/gr.107524.110

27. Pedregosa F, Varoquaux G, Gramfort A, Michel V, Thirion B, Grisel O, et al. Scikit-learn: Machine Learning in Python. Journal of Machine Learning Research. 2011;12: 2825–2830.

28. Lambert SA, Wingfield B, Gibson JT, Gil L, Ramachandran S, Yvon F, et al. Enhancing the Polygenic Score Catalog with tools for score calculation and ancestry normalization. Nature Genetics. 2024;56: 1989–1994.

29. Di Tommaso P, Chatzou M, Floden EW, Barja PP, Palumbo E, Notredame C. Nextflow enables reproducible computational workflows. Nature Biotechnology. 2017;35: 316–319.

